# HIV infection and incidence of cardiovascular diseases: an analysis of a large healthcare database

**DOI:** 10.1101/534024

**Authors:** Alvaro Alonso, A. Elise Barnes, Jodie L. Guest, Amit Shah, Iris Yuefan Shao, Vincent Marconi

## Abstract

**Background:** Persons living with HIV (PLWH) experience higher risk of cardiovascular diseases (CVD) such as myocardial infarction (MI) and heart failure (HF) compared to uninfected individuals. Risk of other CVDs in PLWH has received less attention.

**Methods:** We studied 19,798 PLWH and 59,302 age and sex-matched uninfected individuals identified from the MarketScan Commercial and Medicare databases in the period 2009-2015. Incidence of CVDs, including MI, HF, atrial fibrillation (AF), peripheral artery disease (PAD), stroke and any CVD-related hospitalization, were identified using validated algorithms. We used adjusted Cox models to estimate hazard ratios (HR) and 95% confidence interval (95%CI) of CVD endpoints and performed probabilistic bias analysis to control for unmeasured confounding by race.

**Results:** After a mean (median) follow-up of 20 (17) months, study patients experienced 154 MIs, 223 HF, 93 stroke, 397 AF, 98 PAD and 935 CVD hospitalizations. HR (95%CI) comparing PLWH to uninfected controls were 1.3 (0.9, 1.9) for MI, 3.2 (2.4, 4.2) for HF, 2.7 (1.7, 4.0) for stroke, 1.2 (1.0, 1.5) for AF, 1.1 (0.7, 1.7) for PAD, and 1.7 (1.5, 2.0) for any CVD-related hospitalization. Adjustment for unmeasured confounding led to similar associations [1.2 (0.8, 1.8) for MI, 2.8 (2.0, 3.8) for HF, 2.3 (1.5, 3.6) for stroke, 1.3 (1.0, 1.7) for AF, 0.9 (0.5, 1.4) for PAD, and 1.6 (1.3, 1.9) for CVD hospitalization].

**Conclusion:** Using a large health insurance database, we showed that PLWH have an elevated risk of CVD, particularly HF and stroke. With the aging of the HIV population, developing interventions for cardiovascular health promotion and CVD prevention is imperative.

## INTRODUCTION

The advent and diffusion of combination antiretroviral therapy has led to better survival and reduction of AIDS-related mortality in persons living with HIV (PLWH), with their life expectancy increasing by 9-10 years between 1996 and 2010 in high-income countries.^1^ As a consequence, morbidity and mortality due to aging-related causes among PLWH are on the rise. Currently, over half of the deaths in PLWH in high income countries are non-AIDS related.^2, 3^

Cardiovascular disease (CVD) ranks first as a cause of death in the United States and globally.^4^ Among PLWH, mortality due to CVD is growing in importance. The proportion of deaths attributable to CVD in PLWH in the United States more than doubled between 1999 and 2013, from 2.0% to 4.6%.^5^ Extensive evidence shows that HIV infection is an independent risk factor for coronary artery disease and, to a lesser extent, heart failure (HF).^6, 7^ Information on the link between HIV infection and other cardiovascular conditions such as stroke,^8^ peripheral artery disease (PAD),^9^ or atrial fibrillation (AF),^10^ however, remains scarce. Elucidating the impact of HIV infection on a wide range of CVDs will contribute to a more comprehensive characterization of the burden of non-communicable disorders in PLWH, informing their specific needs for control of CVD risk factors. This is of particular importance given recent findings showing that interventions on traditional cardiovascular risk factors could prevent a substantial proportion of CVD in PLWH.^11^ In addition, this information may provide mechanistic insights into the pathways linking HIV infection and CVD. Thus, we evaluated the association of HIV infection with specific CVDs in a large healthcare administrative database in the United States.

## METHODS

### Study population

We used data from the Truven Health MarketScan Commercial Claims and Encounter Database and the Medicare Supplemental and Coordination of Benefits Database (Truven Health Analytics, Ann Arbor, Michigan) for the period of January 1, 2009 through June 30, 2015. These databases include healthcare claims for inpatient and outpatient services from all levels of care, outpatient pharmacy claims, and enrollment information for individuals enrolled in private health plans or employer-sponsored Medicare supplemental plans across the United States. This study was reviewed and approved by the Institutional Review Board of Emory University. A waiver of the informed consent was obtained.

We identified all patients in the MarketScan databases with an HIV diagnosis (see definition below) and at least 12 months of enrollment before the date of HIV diagnosis. For each HIV case, we selected up to three uninfected individuals matched by sex, year of birth and date of enrollment.

### Definition of HIV status

We defined HIV infection using the presence of any of the following ICD-9-CM codes in any claim: 042 (HIV disease), 079.53 (HIV, type 2), 795.71 (nonspecific serologic evidence of HIV), or V08 (asymptomatic HIV infection status).

### Outcome definition

We defined cardiovascular endpoints applying validated algorithms to diagnostic codes from inpatient and outpatient claims after the date of HIV diagnosis or the corresponding date for uninfected persons. Myocardial infarction (MI) was defined as ICD-9-CM 410 code as primary diagnosis in an inpatient claim,^12^ HF as ICD-9-CM code 428 in any position in an inpatient claim,^13^ stroke as ICD-9-CM codes 430, 431, 434 or 436 as primary diagnosis in an inpatient claim,^14^ AF as ICD-9-CM codes 427.31 or 427.32 in any position in one inpatient claim or two separate outpatient claims,^15^ and PAD as selected ICD-9-CM codes 440, 442, 443 and 444 in any position in an inpatient claim.^16^ Finally, CVD hospitalization was defined as presence of ICD-9-CM codes 390-460 as primary diagnosis code in any inpatient claim. Information on mortality is not available in the MarketScan databases. Supplemental Table S1 includes a detailed list of the codes used for each endpoint.

### Covariates

We used inpatient and outpatient claims up to the time of HIV diagnosis or corresponding date for matched uninfected persons to define potential confounders. We considered comorbidities (hypertension, diabetes, dyslipidemia, coronary artery disease, ischemic stroke, HF, AF, PAD, kidney disease, obesity, sleep apnea, liver disease, thyroid disease, drug abuse, alcohol abuse, smoking) and medication use (lipid lowering medication, antiplatelet, insulin, oral antidiabetics, diuretics, beta blockers, angiotensin receptor blockers, angiotensin converting enzyme inhibitors, non-steroid anti-inflammatory drugs, antidepressants, benzodiazepines). We provide a complete list of ICD-9-CM codes used to define each comorbidity in Supplemental Table S1.

### Statistical analysis

We evaluated the association between HIV infection and risk of each CVD endpoint using multivariable Cox regression, with separate models for each outcome. Time of follow-up was defined as the time between HIV diagnosis, or corresponding date for matched uninfected individuals, and occurrence of the study endpoint, database disenrollment, or September 30, 2015, whichever occurred earlier. An initial model adjusted for age and sex, with an additional analysis including all the covariates defined above. Proportional hazard assumption was explored via inspection of log(-log(survival)) curves and inclusion of time * HIV status terms in the models. We conducted analyses in the overall sample, among those without any prior history of CVD, and stratified by age (<50, ≥50 years) and sex.

MarketScan does not record information on patient race. Not being able to control for race in this analysis is likely to cause confounding given the higher prevalence of HIV infection among blacks in the US and the differences in risk of CVD by race.^17, 18^ To address this limitation, we performed a probabilistic bias analysis to correct for unmeasured confounding following the methodology proposed by Lash and colleagues.^19^ This method corrects the observed estimates of association based on assumptions about the prevalence of the confounder in exposed and unexposed individuals, and the strength of the association between the confounder and the outcomes. To apply this method, we selected a range of values for the proportion of blacks among HIV positive and HIV negative patients, and the strength of the association between race and CVD endpoints. Next, we randomly sampled from this distribution of potential values and corrected the hazard ratio (HR) calculated from the standard analysis using the formula proposed by Schlesselman,^20^ and further correcting for random variability as recommended by Lash et al.^21^ Finally, we repeated the process 10,000 times and calculated bias-corrected HRs and 95% confidence intervals as the median, and 2.5^th^ and 97.5^th^ percentiles of the distribution. We performed this process separately for each CVD endpoint. The online supplement provides details about the methodology, ranges of values for the different parameters (Supplemental Table S2), as well as justification for those values.

## RESULTS

The MarketScan databases included information on 101,545,724 individuals for the period January 1, 2009 through June 30, 2015. Of these, 117,769 had at least one HIV diagnosis claim, with 19,798 of them having at least 12 months of enrollment prior to HIV diagnosis. We selected up to three uninfected persons matched to each HIV infected patient. The final sample size included 79,100 patients (59,302 HIV negative individuals and 19,798 patients with HIV diagnosis) (Figure 1). Table 1 presents patient characteristics by HIV infection status. By design, age and sex distribution in both groups was comparable. In contrast, prevalence of most cardiovascular risk factors, prior history of CVD, and use of several cardiovascular medications was higher in HIV positive individuals than in the uninfected controls.

**Figure 1.**
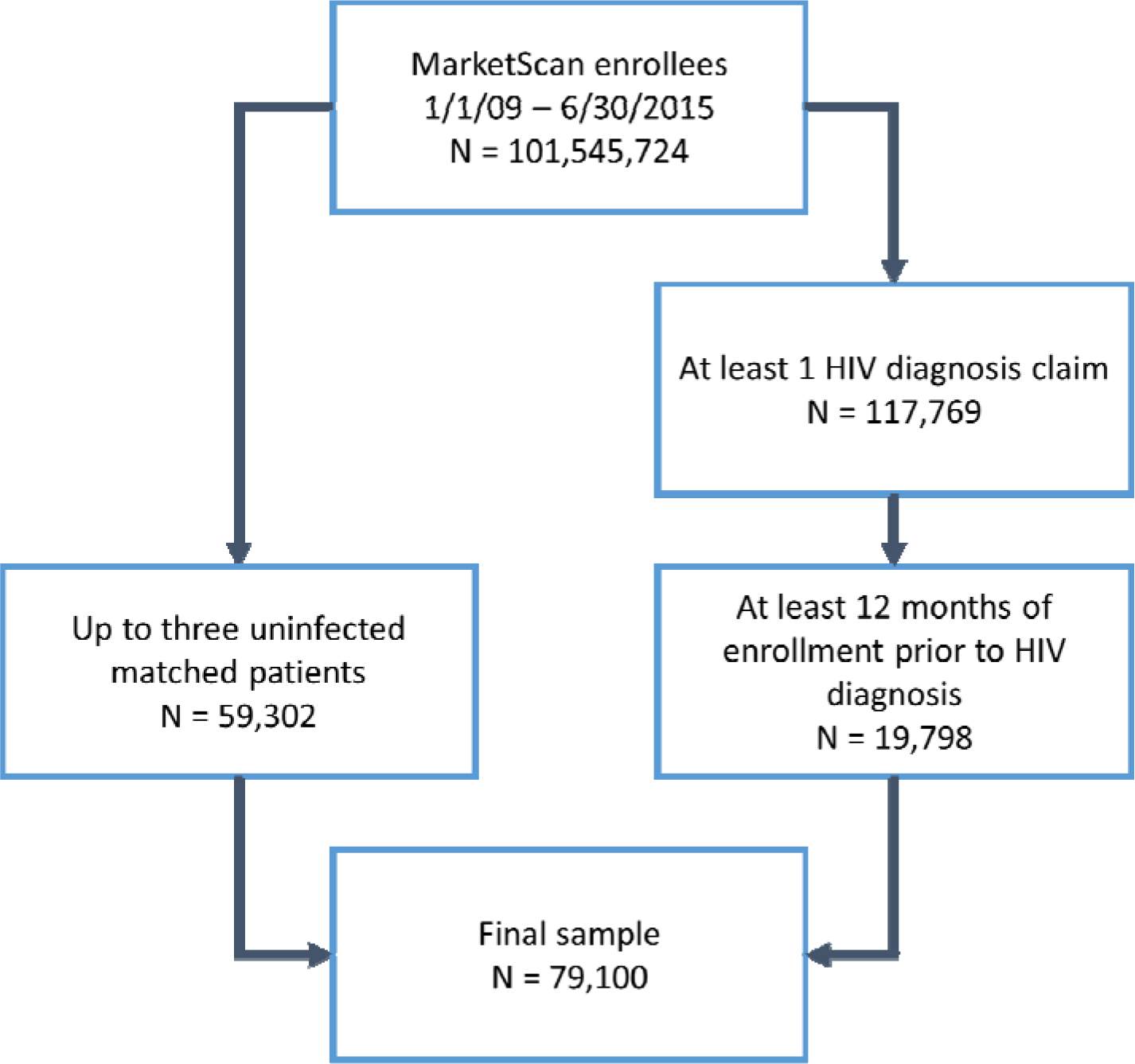
Flowchart of study participants, MarketScan 2009-2015.

**Table 1.**
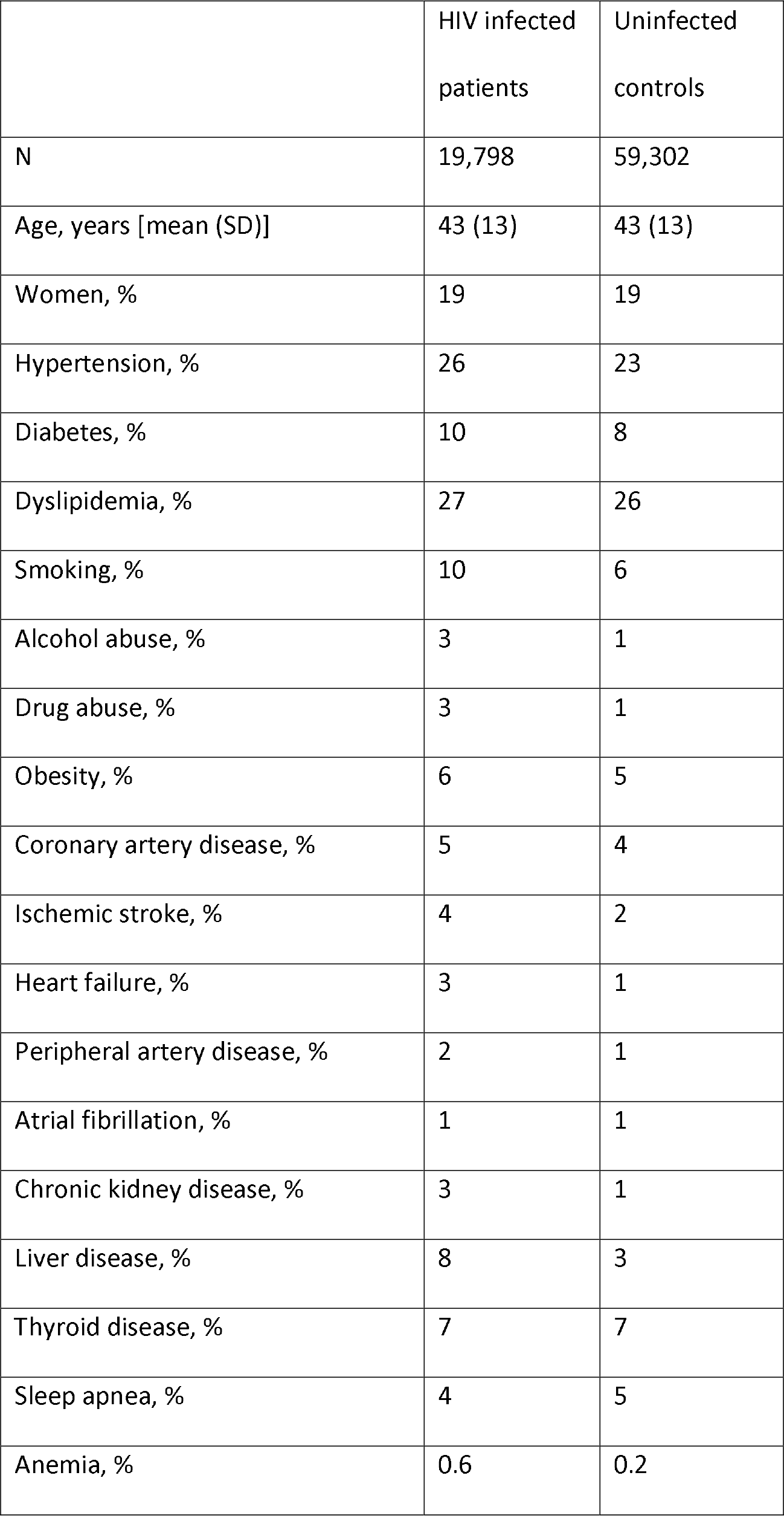
Baseline characteristics by HIV infection status, MarketScan 2009-2015.

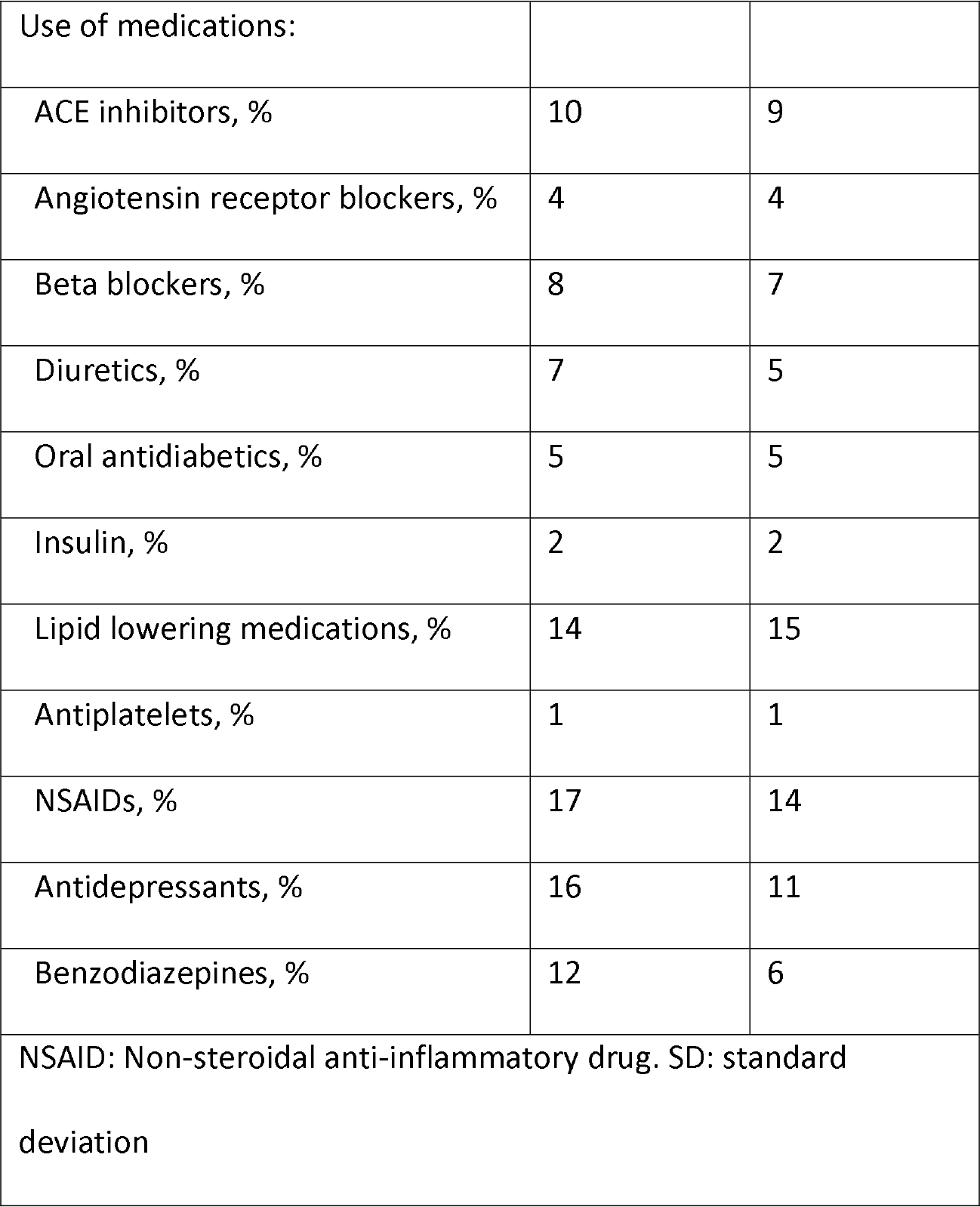

The mean (median) follow-up from index date to disenrollment or September 30, 2015 was 20 (17) months. During follow-up, 935 patients experienced incident cardiovascular hospitalizations, 154 myocardial infarction, 223 heart failure, 98 peripheral artery disease, 93 incident strokes, and 397 atrial fibrillation. Table 2 reports the incident rates of the different cardiovascular events by HIV infection status. The overall rate of CVD hospitalization was 11 per 1,000 person-years among HIV positive and 6 per 1,000 person-years among uninfected individuals. Rates of all individual CVDs were higher among HIV positive than in uninfected individuals.

**Table 2.** Incidence rates of cardiovascular endpoints by HIV infection status, MarketScan 2009-2015.

|  |  | HIV infected<br>patients | Uninfected<br>controls |
| --- | --- | --- | --- |
| CVD hospitalization | N. events | 359 | 576 |
|  | Person-years | 33,394 | 97,449 |
|  | IR (95%CI)* | 10.8 (9.6, 11.9) | 5.9 (5.4, 6.4) |
| Myocardial infarction | N. events | 46 | 108 |
|  | Person-years | 33,873 | 98,135 |
|  | IR (95%CI)* | 1.4 (1.0, 1.8) | 1.1 (0.9, 1.3) |
| Heart failure | N. events | 116 | 107 |
|  | Person-years | 32,939 | 96,897 |
|  | IR (95%CI)* | 3.5 (2.9, 4.2) | 1.1 (0.9, 1.3) |
| Stroke | N. events | 46 | 47 |
|  | Person-years | 33,867 | 98,237 |
|  | IR (95%CI)* | 1.4 (1.0, 1.8) | 0.5 (0.3, 0.6) |
| Peripheral artery disease | N. events | 30 | 68 |
|  | Person-years | 33,304 | 96,942 |
|  | IR (95%CI)* | 0.9 (0.6, 1.2) | 0.7 (0.5, 0.9) |
| Atrial fibrillation | N. events | 116 | 281 |
|  | Person-years | 33,523 | 97,141 |
|  | IR (95%CI)* | 3.5 (2.8, 4.1) | 2.9 (2.6, 3.2) |
| CI: confidence interval. CVD: Cardiovascular disease. IR: Incidence rate |  |  |  |
| * Per 1,000 person-years |  |  |  |

After adjustment for age and sex, rates of CVD hospitalization were double in HIV infected patients compared to uninfected controls, with rates of heart failure and stroke at least tripling in HIV infected patients compared to uninfected controls (Table 3, Model 1). In contrast, rates of myocardial infarction, peripheral artery disease and atrial fibrillation were only 30-40% higher in HIV infected patients than controls. Adjustment for cardiovascular risk factors and other potential confounders had only limited impact in the estimates of association, with HR (95%CI) for heart failure, stroke, and CVD hospitalization of 3.2 (2.4, 4.2), 2.7 (1.7, 4.0), and 1.7 (1.5, 2.0), respectively, in HIV infected patients compared to uninfected individuals (Table 3, Model 2). In contrast, HIV infected patients had no or small increased risk of MI (HR 1.3, 95%CI 0.9, 1.9), AF (HR 1.2, 95%CI 1.0, 1.5) or PAD (HR 1.1, 95%CI 0.7, 1.7) compared to uninfected controls. Associations were of a similar magnitude or slightly stronger when the analysis was restricted to individuals without a prior history of CVD (Supplemental Table S3). Correcting the estimates for unmeasured confounding by race provided similar results, with most associations being slightly attenuated except for that between HIV infection and AF, which was strengthened (Table 3, Model 3).

**Table 3.** Association of HIV infection status with incidence of cardiovascular disease, MarketScan 2009-2015. Values correspond to hazard ratios (95% confidence intervals) comparing HIV infected patients to uninfected controls.

|  | CVD<br>hospitalization | Myocardial<br>infarction | Heart failure | Stroke | Peripheral<br>artery<br>disease | Atrial<br>fibrillation |
| --- | --- | --- | --- | --- | --- | --- |
| Model 1 | 2.0 (1.7, 2.2) | 1.3 (0.9, 1.9) | 3.5 (2.7, 4.6) | 3.0 (2.0, 4.5) | 1.4 (0.9, 2.1) | 1.3 (1.1, 1.6) |
| Model 2 | 1.7 (1.5, 2.0) | 1.3 (0.9, 1.9) | 3.2 (2.4, 4.2) | 2.7 (1.7, 4.0) | 1.1 (0.7, 1.7) | 1.2 (1.0, 1.5) |
| Model 3 | 1.6 (1.3, 1.9) | 1.2 (0.8, 1.8) | 2.8 (2.0, 3.8) | 2.3 (1.5, 3.6) | 0.9 (0.5, 1.4) | 1.3 (1.0, 1.7) |

CVD: Cardiovascular disease
Model 1: Cox proportional hazards model adjusted for age and sex.
Model 2: Cox proportional hazards model adjusted for age, sex, hypertension, diabetes, dyslipidemia, smoking, alcohol abuse (not in incident stroke model), drug abuse (not in incident peripheral artery disease model), obesity, coronary artery disease, ischemic stroke, heart failure (not in incident heart failure model), peripheral artery disease (not in incident peripheral artery disease model), atrial fibrillation (not in incident atrial fibrillation model), chronic kidney disease, liver disease, thyroid disease, sleep apnea, use of ACE inhibitors, angiotensin receptor blockers, beta blockers, diuretics, oral antidiabetics, insulin, lipid lowering medications, antiplatelets, NSAIDs, antidepressants, and benzodiazepines.
Model 3: Results from Model 2 after performing bias analysis for unmeasured confounding by race

The associations were of similar magnitude for men and women, but there was some evidence of effect modification in the HR scale by age. In general, associations were stronger among younger individuals (<50) than older ones (≥50 years of age) (Figure 2). This was particularly notable for HF, with a HR (95%CI) of 5.9 (3.4, 10.1) in the younger group compared to 2.5 (1.8, 3.5) among the older.

**Figure 2.**
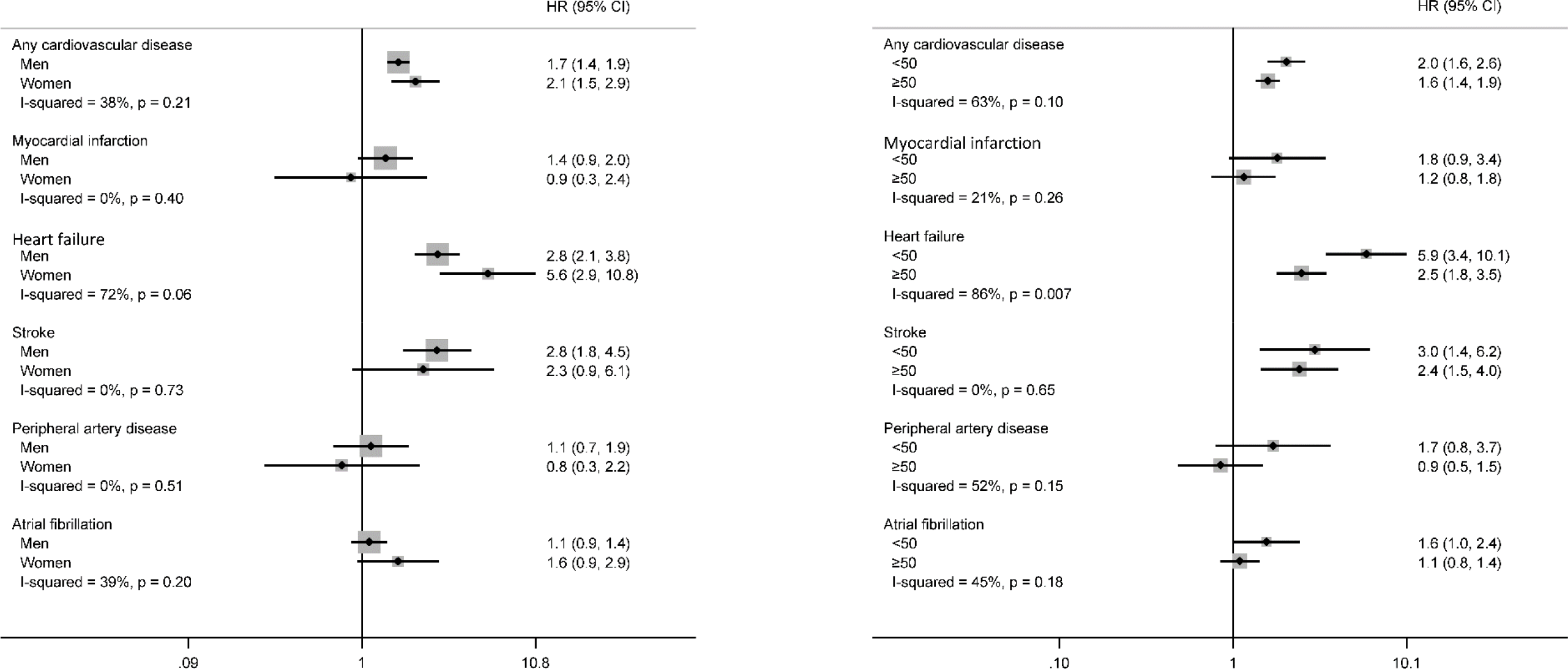
Forest plot presenting associations of HIV infection status with incidence of cardiovascular disease by sex and age group, MarketScan 2009-2015. Results from Cox regression models adjusted for all covariates listed in Model 2 of Table 3.

## DISCUSSION

In this analysis of a large healthcare claims database, we show that PLWH experience higher rates of several CVDs compared with a group of age and sex-matched uninfected individuals, independent of cardiovascular risk factors and other comorbidities. The association of HIV infection with CVD was particularly strong for HF, stroke, and a broadly defined outcome of CVD hospitalization, as well as for persons younger than 50 years of age and those without a prior history of CVD. In this cohort, PLWH did not have increased risk of PAD and only moderately increased risk of MI and AF. Correcting for unmeasured confounding had a minimal impact on the estimates of association.

The extended use of combination antiretroviral therapy has resulted in longer life expectancy in PLWH. Therefore, rates of other chronic conditions, including CVD, in this population have increased. Multiple studies have shown that PLWH are at particularly increased risk of MI and other manifestations of coronary artery disease. For example, the adjusted HR (95%CI) of MI in PLWH in the Veterans Aging Cohort Study Virtual Cohort (mean age 48 years old, 97% men) compared to uninfected controls was 1.48 (1.27, 1.72) in the period 2003-2009.^22^ Our analysis reports a slightly weaker association, consistent with a report from Kaiser Permanente health plans in California suggesting declining relative risks for MI associated with HIV infection in more recent years, from 1.8 (95%CI 1.3, 2.6) in 1996–1999 to 1.0 (95%CI, 0.7, 1.4) in 2010–2011.^23^

The literature regarding CVD endpoints in PLWH other than MI is sparser but growing. In the Veterans Aging Cohort Study, HIV infection was associated with increased risk of both HF with reduced ejection fraction (HR 1.6, 95%CI 1.4, 1.9) and HF with preserved ejection fraction (HR 1.2, 95%CI 1.0, 1.4) compared to no-infection.^24^ Consistent with our findings, associations were stronger in younger than older persons. Similar observations have been made in a cohort of PLWH and uninfected controls in Chicago, with a HR (95%CI) for HF of 2.1 (1.4, 3.2) comparing PLWH to controls.^25^ Direct HIV-induced myocardial damage, myocardial inflammation and fibrosis, and some antiretroviral medications are potential mechanisms underlying this association.^26^ The few studies that have explored an association between HIV infection and stroke reported increased, but relatively weak, rates of stroke among PLWH compared to uninfected individuals: the pooled risk ratios (95%CI) of any stroke and of ischemic stroke in a meta-analysis of published studies were 1.82 (1.53, 2.16) and 1.27 (1.15, 1.39), respectively.^27^ Only one previous publication has reported the prospective association of HIV infection with incidence of PAD. Among 91,953 participants in the Veterans Aging Cohort Study, the HR (95%CI) of PAD was 1.19 (1.13, 1.25) in PLWH versus uninfected persons.^9^ These results are consistent with the findings from our analysis. Finally, no published studies have compared directly rates of AF in PLWH and uninfected persons. An analysis of 30,533 PLWH from the Veterans Affairs HIV Clinical Case Registry reported an incidence of AF of 3.6 events per 1,000 person-years, but no uninfected comparison group was included.^10^

The present study makes some significant contributions to the extensive literature on the relationship between HIV infection and incidence of CVD. First, it shows that healthcare claims databases are a valuable resource for the epidemiologic study of CVD and other chronic conditions in PLWH. Carefully phenotyped prospective cohorts of PLWH and uninfected persons are uniquely suited to answer questions about outcomes associated with HIV infection, but they can be limited by their sample size, geographic restrictions, lack of diversity, or narrow list of disease endpoints. Large claims databases, despite their known shortcomings, provide large sample size, include patients from wide geographic areas, and reflect the sociodemographic diversity of the general insured population. Second, ours is the first study to compare incidence of AF in PLWH to an uninfected cohort. We report that PLWH had a 40% increased risk of AF compared to uninfected individuals, independent of other cardiovascular risk factors and taking into account unmeasured confounding. Inflammation associated with HIV infection as well as direct effects of HIV on the atrial myocardium, as shown by studies reporting high prevalence of left atrial enlargement and myocardial fibrosis in PLWH,^28, 29^ could explain this association. A previous study indicating that markers of HIV infection severity were associated with incident of AF among PLWH indirectly supports a deleterious effect of HIV on AF pathogenesis.^10^ Third, we assessed a wide range of CVD endpoints, providing a comprehensive characterization of the cardiovascular risk in this population, and highlighting areas that require particular attention beyond the prevention of coronary artery disease, such as the high rate of HF hospitalization among PLWH.

We used bias analysis to evaluate the impact of unmeasured confounding by race in our estimates of association. Risk of HIV and most CVDs is higher among African Americans compared to other racial/ethnic groups in the United States, particularly whites.^17, 30^ Unfortunately, the MarketScan database does not include information on race, restricting our ability to control for this variable. Making reasonable assumptions about the proportion of blacks in PLWH and uninfected persons and the association of race with the different CVD endpoints, and applying well-established methods, we were able to obtain estimates of association adjusting for unmeasured confounding by race. As expected, estimates for most CVDs were attenuated in the corrected analysis. The only exception was AF, where the association between HIV status and AF risk became stronger. This is expected since, in contrast to other CVDs, rates of AF are lower in African Americans than whites,^31^ and confounding by race would attenuate the true association.

The present analysis has some important strengths including the large patient population, the use of validated algorithms to define CVD endpoints, the ability to control for a variety of confounders and the use of bias analysis to mitigate the threat of unmeasured confounding. However, certain limitations must be highlighted. First, we relied on claims to define exposures, outcomes and covariates, which can lead to misclassification even when using validated algorithms. Lack of information on out-of-hospital fatal events also causes outcome misclassification. Second, increased healthcare utilization among PLWH may contribute to increased ascertainment of CVD outcomes, which could lead to an apparent higher risk of CVD in the HIV infected group. Third, claims are inadequate to evaluate HIV infection severity, since information such as viral load, CD4+ count or compliance with prescribed antiretroviral therapy is not available. Fourth, despite adjusting for numerous covariates and using bias analysis, uncontrolled confounding is likely to remain, including psychological and social risk factors more prevalent among PLWH. Fifth, our study sample is restricted to individuals with commercial or Medicare supplemental insurance in the United States and, thus, results may not generalize to the uninsured, those in other type of insurance plans (e.g. Medicaid), or to persons outside the United States.

To conclude, we observed that PLWH are at increased risk of several CVDs. Results from this study can inform future research by highlighting conditions that require increased attention and providing clues regarding the mechanisms that put PLWH at greater risk for certain CVDs compared to their uninfected counterparts. More work is needed to help with recognition, prevention and treatment of CVD among PLWH.

## Supporting information

Supplemental Methods and Results

## REFERENCES

1. Antiretroviral Therapy Cohort Collaboration. Survival of HIV-positive patients starting antiretroviral therapy between 1996 and 2013: a collaborative analysis of cohort studies. Lancet HIV. 2017;4:e349–e356.

2. Antiretroviral Therapy Cohort Collaboration. Causes of death in HIV-1-infected patients treated with antiretroviral therapy, 1996-2006: collaborative analysis of 13 HIV cohort studies. Clin Infect Dis. 2010;50:1387–1396.

3. Farahani M, Mulinder H, Farahani A, Marlink R. Prevalence and distribution of non-AIDS causes of death among HIV-infected individuals receiving antiretroviral therapy: a systematic review and meta-analysis. Int J STD AIDS. 2017;28:636–650.

4. GBD 2016 Causes of Death Collaborators. Global, regional, and national age-sex specific mortality for 264 causes of death, 1980-2016: a systematic analysis for the Global Burden of Disease Study 2016. Lancet. 2017;390:1151–1210.

5. Feinstein MJ, Bahiru E, Achenbach C, et al. Patterns of cardiovascular mortality for HIV-infected adults in the United States: 1999 to 2013. Am J Cardiol. 2016;117:214–220.

6. Kaplan RC, Hanna DB, Kizer JR. Recent insights into cardiovascular disease (CVD) risk among HIV-infected adults. Curr HIV/AIDS Rep. 2016;13:44–52.

7. So-Armah K, Freiberg MS. HIV and cardiovascular disease: update on clinical events, special populations, and novel biomarkers. Curr HIV/AIDS Rep. 2018;doi: 10.1007/s11904-018-0400-5. [Epub ahead of print].

8. Sico JJ, Chang CC, So-Armah K, et al. HIV status and the risk of ischemic stroke among men. Neurology. 2015;84:1933–1940.

9. Beckman JA, Duncan MS, Alcorn CW, et al. Association of HIV infection and risk of peripheral artery disease. Circulation. 2018;138:255–265.

10. Hsu JC, Li Y, Marcus GM, et al. Atrial fibrillation and atrial flutter in human immunodeficiency virus-infected persons: incidence, risk factors, and association with marker of HIV disease severity. J Am Coll Cardiol. 2013;61:2288–2295.

11. Althoff KN, Gebo KA, Moore RD, et al. Contributions of traditional and HIV-related risk factors on non-AIDS-defining cancer, myocardial infarction, and end-stage liver and renal diseases in adults with HIV in the USA and Canada: a collaboration of cohort studies. Lancet HIV. 2019;in press. DOI: 10.1016/S2352-3018(18)30295-9.

12. Cutrona SL, Toh S, Iyer A, et al. Validation of acute myocardial infarction in the Food and Drug Administration’s Mini-Sentinel program. Pharmacoepidemiol Drug Saf. 2013;22:40–54.

13. Loehr LR, Rosamond WD, Chang PP, Folsom AR, Chambless LE. Heart failure incidence and survival (from the Atherosclerosis Risk in Communities Study). Am J Cardiol. 2008;101:1016–1022.

14. Andrade SE, Harrold LR, Tjia J, et al. A systematic review of validated methods for identifying cerebrovascular accident or transient ischemic attack using administrative data. Pharmacoepidemiol Drug Saf. 2012;21 Supplement 1:100–128.

15. Jensen PN, Johnson K, Floyd J, Heckbert SR, Carnahan R, Dublin S. A systematic review of validated methods for identifying atrial fibrillation using administrative data. Pharmacoepidemiol Drug Saf. 2012;21 Suppl 1:141–147.

16. Hirsch AT, Hartman L, Town RJ, Virnig BA. National health costs of peripheral arterial disease in the Medicare population. Vasc Med. 2008;13:209–215.

17. Centers for Disease Control and Prevention. HIV Surveillance Report, 2017 2018.

18. Rosamond WD, Chambless LE, Heiss G, et al. Twenty-two-year trends in incidence of myocardial infarction, coronary heart disease mortality, and case fatality in 4 US communities, 1987-2008. Circulation. 2012;125:1848–1857.

19. Lash TL, Fox MP, Fink AK. Probabilistic bias analysis. Applying quantitative bias analysis to epidemiologic data. New York: Springer; 2009:117–150.

20. Schlesselman JJ. Assessing effects of confounding variables. Am J Epidemiol. 1978;108:3–8.

21. Lash TL, Fox MP, Fink AK. Applying quantitative bias analysis to epidemiologic data. New York: Springer; 2009.

22. Freiberg MS, Chang CC, Kuller LH, et al. HIV infection and the risk of acute myocardial infarction. JAMA Intern Med. 2013;173:614–622.

23. Klein DB, Leyden WA, Xu L, et al. Declining relative risk for myocardial infarction among HIV-positive compared with HIV-negative individuals with access to care. Clin Infect Dis. 2015;60:1278–1280.

24. Freiberg MS, Chang CH, Skanderson M, et al. Association between HIV infection and the risk of heart failure with reduced ejection fraction and preserved ejection fraction in the antiretroviral therapy era: results from the Veterans Aging Cohort Study. JAMA Cardiol. 2017;2:536–546.

25. Feinstein MJ, Stevenson AB, Ning H, et al. Adjudicated heart failure in HIV-infected and uninfected men and women. J Am Heart Assoc. 2018;7:e009985.

26. Savvoulidis P, Butler J, Kalogeropoulos A. Cardiomyopathy and heart failure in patients with HIV infection. Can J Cardiol. 2018;(in press) DOI: 10.1016/j.cjca.2018.10.009.

27. Gutierrez J, Albuquerque ALA, Falzon L. HIV infection as vascular risk: a systematic review of the literature and meta-analysis. PLoS One. 2017;12:e0176686.

28. Mondy KE, Gottdiener J, Overton ET, et al. High prevalence of echocardiographic abnormalities among HIV-infected persons in the era of highly active antiretroviral therapy. Clin Infect Dis. 2011;52:378–386.

29. Thiara DK, Liu CY, Raman F, et al. Abnormal myocardial function is related to myocardial steatosis and diffuse myocardial fibrosis in HIV-infected adults. J Infect Dis. 2015;212:1544–1551.

30. Benjamin EJ, Virani SS, Callaway CW, et al. Heart disease and stroke statistics—2018 update: a report from the American Heart Association. Circulation. 2018;137:e67–e492.

31. Alonso A, Agarwal SK, Soliman EZ, et al. Incidence of atrial fibrillation in whites and African-Americans: the Atherosclerosis Risk in Communities (ARIC) study. Am Heart J. 2009;158:111–117.

