## Supplemental Methods and Results for "HIV infection and incidence of cardiovascular diseases: an analysis of a large healthcare database"

**SUPPLEMENTAL MATERIAL**

**Supplemental methods**

Information on race is not available in the MarketScan databases. Given the known association of race with HIV infection and CVD risk in the United States, our analyses in the MarketScan database without race adjustment are hopelessly confounded. To address this issue, we corrected our estimates of association using probabilistic bias analysis. This approach calculates adjusted estimates of association using the observed associations, the prevalence of the confounder in the exposed and unexposed, and the strength of the association between the confounder and the outcome of interest. We followed the approach recommended by Lash and colleagues to perform this analysis:

1. In a first step, we assigned a distribution for bias analysis parameters based on published literature (detailed in Supplemental Table S2). For most parameters, we used a trapezoidal distribution, with minimum and maximum defined as the limits of 95% confidence intervals for estimates in published studies, and lower and upper modes based on point estimates in those publications.
2. Second, we randomly sampled parameters from that distribution and corrected the observed measure of association using the following formula:

$${HR}_{adj}={HR}_{observed}\frac{{HR}_{CD}p_{0}+\left( 1-p_{0} \right)}{{HR}_{CD}p_{1}+\left( 1-p_{1} \right)}$$

where *HR*_adj_ is the hazard ratio for the association of the exposure and the endpoint adjusted for the unmeasured confounder, *HR*_observed_ is the observed hazard ratio for that association, *HR_CD_* is the hazard ratio for the association of the confounder with the disease, and *p*_1_ and *p*_0_ are the proportion of subjects with the confounder in the exposed and unexposed, respectively.^1^ This formula assumes that there is no effect measure modification of the confounder-disease association by the exposure, and that the HR is an adequate estimate of the risk ratio.

3. Third, we added random error to the bias-corrected associations using the following formula:

$${HR}_{total}=exp\left[ \ln\left( {HR}_{systematic} \right)-{random}_{0,1}\cdot{ste}_{conventional}) \right]$$

Where *HR*_total_ is the estimate of association incorporating both systematic and random error, *HR*_systematic_ is the estimate that corrects for unmeasured confounding, *random*_0,1_ is a random number from a normal distribution of mean equals 0 and standard deviation equals 1, and *ste*_conventional_ is the standard error of the conventional estimate of association.^2^

4. Finally, we repeated this process 10,000 times, and calculated the final estimate of association as the median of the distribution of the 10,000 estimates, with a 95% confidence interval using the 2.5^th^ and the 97.5^th^ percentiles.

**Supplemental Table S1**. Diagnosis codes used to define endpoints or covariates in the study population.

| **Condition** | **ICD-9-CM codes** |
| --- | --- |
| *Cardiovascular endpoints* |  |
| Myocardial infarction | 410.xx as primary diagnosis in inpatient claim |
| Heart failure | 428.xx in any position in inpatient claim |
| Stroke | 430.xx, 431.xx, 434.xx, 436.xx as primary diagnosis in inpatient claim |
| Atrial fibrillation | 427.3x in any position in 1 inpatient or 2 outpatient claims |
| Peripheral artery disease | 440.0x, 440.20, 440.21, 440.22, 440.23, 440.24, 440.31, 440.9x, 442.3x, 443.9x, 444.2x, 444.81 in any position in inpatient claim |
| Any cardiovascular disease hospitalization | 390-460 as primary diagnosis in inpatient claim |
| *Comorbidities** |  |
| Hypertension | 401.xx, 402.xx, 403.xx, 404.xx, 405.xx |
| Diabetes | 250.xx |
| Dyslipidemia | 272.xx |
| Smoking | 305.1, 649.0x, 989.84, V15.82 |
| Coronary artery disease | 410.xx, 411.xx, 412.xx, 413.xx, 414.xx |
| Ischemic stroke | 434.xx, 436.xx |
| Obesity | 278.0x |
| Sleep apnea | 327.2x, 780.51, 780.53, 780.57 |
| Chronic kidney disease | 403.01, 403.11, 403.91, 404.02, 404.03, 404.12, 404.13, 404.92, 404.93, 582.xx, 583.0x, 583.1x, 583.2x, 583.3x, 583.4x, 583.5x, 583.6x, 583.7x, 585.xx, 586.xx, 588.0x, V42.0x, V45.1x, V56.xx |
| Liver disease | 070.22, 070.23, 070.32, 070.33, 070.44, 070.54, 070.6x, 070.9x, 456.0x, 456.1x, 456.2x, 570.xx, 571.xx, 572.2x, 572.3x, 572.4x, 572.5x, 572.6x, 572.7x, 572.8x, 573.3x, 573.4x, 573.8x, 573.9x, V42.7 |
| Thyroid disease | 240.xx, 241.xx, 242.xx, 244.xx, 245.xx, 246.xx |
| Drug abuse | 292.xx, 304.xx, 305.2x, 305.3x, 305.4x, 305.5x, 305.6x, 305.7x, 305.8x, 305.9x |
| Anemia | 290.xx, 291.xx |
| Alcohol abuse | 265.2x, 291.1x, 291.2x, 291.3x, 291.5x, 291.6x, 291.7x, 291.8x, 291.9x, 303.0x, 303.9x, 305.0x, 357.5x, 425.5x, 535.3x, 571.0x, 571.1x, 571.2x, 571.3x, 980.xx, V11.3 |
| * Comorbidities were defined based on the presence of the indicated codes in any position in inpatient or outpatient claims. ICD-9-CM: International Classification of Diseases, 9^th^ revision, Clinical Modification | |

**Supplemental Table S2**. Bias analysis parameters for race as an unmeasured confounder. References for the sources of estimates are indicated in the table.

|  | Minimum | Lower mode | Upper mode | Maximum |
| --- | --- | --- | --- | --- |
| Prevalence of black race by HIV status | | | | |
| HIV negative, %^3^ | 0 | 3 | 15 | 38 |
| HIV positive, %^4^ | 16 | 38 | 43 | 54 |
| Hazard ratios of black race and CVD outcome | | | | |
| Any CVD^5^ | 1.39 | 1.45 | 1.45 | 1.51 |
| MI^6^ | 0.94 | 1.15 | 1.48 | 1.90 |
| HF^7, 8^ | 1.39 | 1.55 | 1.80 | 2.00 |
| Stroke^9, 10^ | 1.26 | 1.51 | 2.03 | 2.30 |
| PAD^11, 12^ | 1.36 | 1.41 | 2.33 | 3.99 |
| AF^13, 14^ | 0.38 | 0.51 | 0.59 | 0.92 |

**Supplemental Table S3**. Association of HIV infection status with incidence of cardiovascular disease among individuals without any history of cardiovascular disease, MarketScan 2009-2015. Values correspond to hazard ratios (95% confidence intervals) comparing HIV positive to HIV negative patients.

|  | HIV positive (N = 17,843) | | HIV negative (N = 55,281) | |  |
| --- | --- | --- | --- | --- | --- |
|  | N. events | Person-years | N. events | Person-years | HR (95%CI)^a^ |
| CVD hospitalization | 217 | 30,471 | 331 | 90,879 | 2.1 (1.8, 2.5) |
| Myocardial infarction | 28 | 30,783 | 75 | 91,290 | 1.3 (0.8, 2.0) |
| Heart failure | 85 | 30,714 | 66 | 91,320 | 3.8 (2.8, 5.3) |
| Stroke | 36 | 30,777 | 29 | 91,354 | 3.7 (2.2, 6.1) |
| Peripheral artery disease | 22 | 30,760 | 41 | 91,364 | 1.5 (0.9, 2.6) |
| Atrial fibrillation | 89 | 30,709 | 189 | 91,151 | 1.5 (1.2, 2.0) |

CI: confidence interval. CVD: cardiovascular disease. HR: hazard ratio

^a^Cox proportional hazards model adjusted for age, sex, hypertension, diabetes, dyslipidemia, smoking, alcohol abuse (not in incident stroke model), drug abuse (not in incident peripheral artery disease model), obesity, chronic kidney disease, liver disease, thyroid disease, sleep apnea, use of ACE inhibitors, angiotensin receptor blockers, beta blockers, diuretics, oral antidiabetics, insulin, lipid lowering medications, antiplatelets, NSAIDs, antidepressants, and benzodiazepines.

**SUPPLEMENTAL REFERENCES**

**1.**	Schlesselman JJ. Assessing effects of confounding variables. *Am J Epidemiol.* 1978;108:3-8.

**2.**	Lash TL, Fox MP, Fink AK. *Applying quantitative bias analysis to epidemiologic data*. New York: Springer; 2009.

**3.**	American Community Survey. In: US Census Bureau, ed; 2017.

**4.**	Centers for Disease Control and Prevention. *HIV Surveillance Report, 2017* 2018.

**5.**	Oramasionwu CU, Morse GD, Lawson KA, Brown CM, Koeller JM, Frei CR. Hospitalizations for cardiovascular disease in African Americans and whites with HIV/AIDS. *Popul Health Manag.* 2013;16:201-207.

**6.**	Safford MM, Brown TM, Muntner PM, et al. Association of race and sex with risk of incident acute coronary heart disease events. *JAMA.* 2012;308:1768-1774.

**7.**	Loehr LR, Rosamond WD, Chang PP, Folsom AR, Chambless LE. Heart failure incidence and survival (from the Atherosclerosis Risk in Communities Study). *Am J Cardiol.* 2008;101:1016-1022.

**8.**	Chang PP, Wruck LM, Shahar E, et al. Trends in hospitalizations and survival of acute decompensated heart failure in four US communities (2005-2014): ARIC Study Community Surveillance. *Circulation.* 2018;138:12-24.

**9.**	Howard VJ, Kleindorfer DO, Judd SE, et al. Disparities in stroke incidence contributing to disparities in stroke mortality. *Ann Neurol.* 2011;69:619-627.

**10.**	Koton S, Schneider AL, Rosamond WD, et al. Stroke incidence and mortality trends in US communities, 1987 to 2011. *JAMA.* 2014;312:259-268.

**11.**	Kalbaugh CA, Kucharska-Newton A, Wruck L, et al. Peripheral artery disease prevalence and incidence estimated from both outpatient and inpatient settings among Medicare fee-for-service beneficiaries in the Atherosclerosis Risk in Communities (ARIC) Study. *J Am Heart Assoc.* 2017;6:e003796.

**12.**	Vart P, Coresh J, Kwak L, Ballew SH, Heiss G, Matsushita K. Socioeconomic status and incidence of hospitalization with lower-extremity peripheral artery disease: Atherosclerosis Risk in Communities Study. *J Am Heart Assoc.* 2017;6:e004995.

**13.**	Alonso A, Agarwal SK, Soliman EZ, et al. Incidence of atrial fibrillation in whites and African-Americans: the Atherosclerosis Risk in Communities (ARIC) study. *Am Heart J.* 2009;158:111-117.

**14.**	Rodriguez CJ, Soliman EZ, Alonso A, et al. Atrial fibrillation incidence and risk factors in relation to race-ethnicity and the population attributable fraction of atrial fibrillation risk factors: the Multi-Ethnic Study of Atherosclerosis. *Ann Epidemiol.* 2015;25:71-77.
